# Genomic streamlining of seagrass-associated *Colletotrichum* sp. may be related to its adaptation to a marine monocot host

**DOI:** 10.1101/2024.12.17.629027

**Authors:** Cassandra L. Ettinger, Jonathan A. Eisen, Jason E. Stajich

## Abstract

*Colletotrichum* spp. have a complicated history of association with land plants. Perhaps most well-known as plant pathogens for the devastating effect they can have on agricultural crops, some *Colletotrichum* spp. have been reported as beneficial plant endophytes. However, there have been only a handful of reports of *Colletotrichum* spp. isolated from aquatic plant hosts and their ecological role in the marine ecosystem is underexplored. To address this, we present the draft genome and annotation of *Colletotrichum* sp. CLE4, previously isolated from rhizome tissue from the seagrass *Zostera marina*. This genome (48.03 Mbp in length) is highly complete (BUSCO ascomycota: 98.8%) and encodes 12,015 genes, of which 5.7% are carbohydrate-active enzymes (CAZymes) and 12.6% are predicted secreted proteins. Phylogenetic placement puts *Colletotrichum* sp. CLE4 within the *C. acutatum* complex, closely related to *C. godetiae*. We found a 8.69% smaller genome size, 21.90% smaller gene count, and the absence of 591 conserved gene families in *Colletotrichum* sp. CLE4 relative to other members of the *C. acutatum* complex, suggesting a streamlined genome possibly linked to its specialized ecological niche in the marine ecosystem. Machine learning analyses using CATAStrophy on CAZyme domains predict this isolate to be a hemibiotroph, such that it has a biotrophic phase where the plant is kept alive during optimal environmental conditions followed by a necrotrophic phase where the fungi actively serves a pathogen. While future work is still needed to definitively tease apart the lifestyle strategy of *Colletotrichum* sp. CLE4, this study provides foundational insight and a high-quality genomic resource for starting to understand the evolutionary trajectory and ecological adaptations of marine-plant associated fungi.

## Introduction

*Colletotrichum* is a diverse genus of plant-associated fungi well known as both pathogens and endophytes of terrestrial plants [1]. Many *Colleotrichum* exhibit a complex hemibiotrophic lifestyle, meaning they have an initial biotrophic phase where the plant host is kept alive, followed by a necrotrophic phase where the fungi actively harms host tissues [2]. During this necrotrophic phase, *Colleotrichum* spp. cause a significant number of diseases, known as anthracnose, in many agricultural crops worldwide, and thus has been named one of the ten most important fungal pathogens [3]. As a result of their complex lifestyle *Colleotrichum* spp. are highly adaptable, associating with a large host range of over 3,200 species of monocot and eudicot plants [4]. While some *Colleotrichum* species have high host specificity, including one-to-one associations, others can infect a wide variety of plant hosts [2,4–7]. Evolutionary analyses suggest that the ancestor of *Colletotrichum* diverged in parallel with the diversification of flowering plants on land, likely beginning with an association with eudicot plants before adapting to other host types [6].

While *Colletotrichum* spp. are predominantly known for their associations with land plants, there have been recent reports documenting their presence as endophytes of aquatic plants [8–10]. Notably, *Colletotrichum* species have been isolated as endophytes in seagrasses, including the ecologically important species *Zostera marina* [11]. *Z. marina* is an early diverging marine monocot that serves as a foundation species in coastal ecosystems across the Northern Hemisphere, with critical roles providing habitat, stabilizing sediment, and contributing to carbon sequestration [12–14].

Previous amplicon-based surveys and culture-dependent studies have reported that *Colletotrichum* spp. are abundant members of the fungal community associated with *Z. marina*, particularly on leaf tissues [11,15]. These studies have found *Colletotrichum* spp. to be dominant on and within healthy *Z. marina* leaves and present in rhizomes, suggesting a possible endophytic relationship. Furthermore, global mycobiome surveys of *Z. marina* have predicted *Colletotrichum* spp. members to be dispersal-limited and exhibit patterns of endemism to specific locations, such as California and Japan [16]. Additionally, *Colletotrichum* spp. have been isolated from leaves and rhizomes of another seagrass species, *Thalassia testudinum*, further supporting their potential role as endophytes in marine environments [17,18].

Given the pathogenic potential of *Colletotrichum* spp. in terrestrial plants, where they cause black lesions characteristic of anthracnose, it is crucial to understand their ecological role in seagrass environments. While no true fungi have yet been reported to cause widespread disease in seagrasses [19], *Colletotrichum* spp. lesions might appear morphologically similar to and be mistaken for those caused by the heterokont pathogen *Labyrinthula zosterae*, which is responsible for seagrass wasting disease [20]. As climate change continues to impact marine ecosystems, it is critical to understand the role of fungi like *Colletotrichum* sp. CLE4 in seagrass health and disease dynamics.

To start to investigate the ecology of seagrass-associated *Colletotrichum* species, we generated a draft genome and annotation for *Colletotrichum* sp. CLE4, previously isolated as an endophyte from the seagrass *Zostera marina* in Ettinger & Eisen [11]. We used this genome to refine taxonomic understanding of this isolate through whole-genome phylogenetic placement among close relatives. We further conducted comparative genomic analyses to identify genes that might have been gained or lost during adaptation to a marine monocot host, leveraging the genome annotation to explore potential ecological roles of *Colletotrichum* sp. CLE4 in the marine environment.

## Methods

### Molecular methods

*Colletotrichum* sp. CLE4 was previously isolated from healthy *Z. marina* rhizome tissues collected in May 2018 from Bodega Bay, CA using Potato Dextrose Agar with 0.45 μM Millipore filtered natural aged seawater as described in Ettinger & Eisen [11] (Figure 1A). Briefly, in that work, the isolate was propagated on the same solid media and DNA was extracted from tissue using a MoBio PowerSoil DNA Extraction kit. The isolate was then identified through phylogenetic analysis using ITS-LSU regions obtained through Sanger sequencing (GenBank Accession: MN543905). In this work, that same DNA was provided to the UC Davis Genome Center DNA Technologies Core for genomic library preparation and sequencing. DNA libraries were sequenced on an Illumina HiSeq4000 to generate 150 bp paired-end reads.

**Figure 1.**
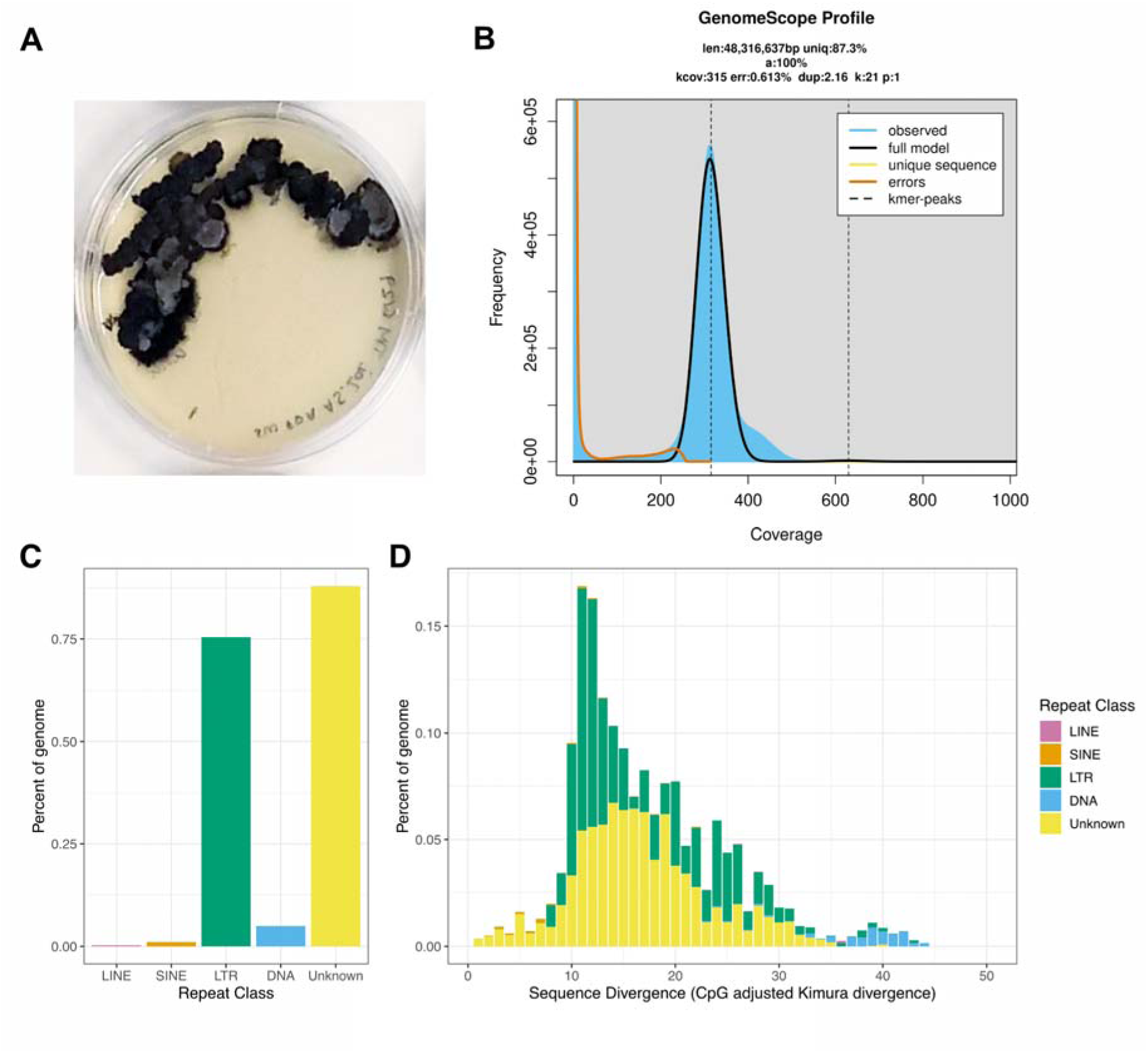
General characteristics of *Colletotrichum* sp. CLE4. (A) Photograph of *Colletotrichum* sp. CLE4. (B) GenomeScope profile depicting the *k-*mer frequency histogram used to calculate genome size, ploidy and heterozygosity. (C) A bar plot representing the percent of the genome composed of repetitive elements from each repeat class. (D) A stacked bar plot representing the percent of the genome made of repeat elements from each repeat class binned by 1% sequence divergence (CpG adjusted Kimura divergence). For (C) and (D) bars are colored repeat class (LINE = pink, SINE = orange, LTR = green, DNA = light blue, and Unknown = yellow). Abbreviations: long-interspersed nuclear element (LINE), small-interspersed nuclear element (SINE), long-terminal repeat retrotransposon (LTR), and DNA transposons (DNA).

### Assembly and annotation

Reads were assembled using the Automatic Assembly of the Fungi (AAFTF) pipeline v. 0.2.5 [21]. This pipeline trims and filters reads using BBTools v. 38.95 [22]. Then, AAFTF assembles these trimmed reads with SPAdes v. 3.14.1 [23] using default parameters. AAFTF screens the resulting assembly for contaminant vectors using BLAST and then uses sourmash v. 3.5.0 [24] to identify and remove any additional contaminant contigs. AAFTF then identifies duplicate contigs for removal using Minimap2 v. 2.17 [25]. Finally, AAFTF runs Pilon v. 1.22 [26] with three rounds of polishing to produce short-read corrected contigs in the assembly.

Repetitive regions were identified and masked prior to genome annotation using RepeatModeler v. 2.0.1 [27] and RepeatMasker v. 4-1-1 [28] with default options to produce a *de novo* library of elements plus the elements from Fungi in Repbase to mask [29]. The repeat landscape of *Colletotrichum* sp. CLE4 was visualized in R v. 4.3.0 [30] using the tidyverse v. 2.0.0 [31]. We then used the Funannotate pipeline v. 1.8.8 to annotate the draft *Colletotrichum* sp. CLE4 genome assembly [32]. Funannotate uses a combination of software to predict gene models including Augustus v. 3.3.3, GlimmerHMM v. 3.0.4, GeneMark-ETS v. 4.62, and SNAP v. 2013_11_29 [33–37], and produces consensus gene model predictions using EVidenceModeler v. 1.1.1 [38]. Funannotate additionally predicts tRNAs using tRNAscan v. 1.3.1 [39]. Funannotate then annotates consensus gene models based on similarity to Pfam-A v. 35.0 [40] and dbCAN v. 9.0 [41,42] using HMMER v.3 [43] and similarity to MEROPS v. 12.0 [44], eggNOG v. 2.1.9 [45], InterProScan v. 5.51-85.0 [46], and UniProt v. 2022_05 [47] using diamond BLASTP v. 2.0.8 [48]. Additionally, Funannotate uses Phobius v. 1.01 [49] to predict transmembrane proteins and SignalP v. 5.0b [50] to predict secreted proteins. AntiSMASH v. 6.1.1 was used to further identify biosynthetic gene clusters [51]. EffectorP v. 3.0 was run on predicted secreted proteins to predict plant effectors [52].

The draft assembly and predicted gene models were assessed for completion using BUSCO v. 5.0.0 [53] in ‘genome’ and ‘protein’ mode, respectively, with the eukartyota_odb10, fungi_odb10 and ascomycota_odb10 sets. To assess genome size and ploidy, we used jellyfish v. 2.3.0 [54] with a *k*-mer size of 21 to produce a *k*-mer frequency histogram, which we supplied to GenomeScope v. 2.0 [55] to predict haploid genome size and heterozygosity.

We used CATAStrophy v. 0.1.0 [56], a classification method based on carbohydrate-active enzyme (CAZyme) patterns from filamentous fungal plant pathogens, to predict the possible lifestyle strategy of *Colletotrichum* sp. CLE4. CATAStrophy was run in a Google Collab implementation using dbCAN v. 10 [41,42].

### Comparative genomics

Predicted gene models from annotated genomes were downloaded from either NCBI or JGI for use in comparative analyses. Completion of downloaded predicted gene models was assessed using BUSCO v. 5.0.0 in ‘protein’ mode with the eukartyota_odb10, fungi_odb10 and ascomycota_odb10 sets. To be included in analysis, the annotated protein sets needed to have >90% completion. In total the final dataset represented 110 annotated genomes [6,57–99] (Table S1). Briefly in addition to *Colletotrichum* sp. CLE4, the dataset included an in-group of 60 *Colletotrichum* genomes and an outgroup representing 49 genomes across six taxonomic orders including Diaporthales (n=2), Glomerellales (n=2), Hypocreales (n=18), Ophiostomatales (n=2), Sordariales (n=5), and Xylariales (n=20). Downloaded gene models were annotated using InterProScan v. 5.51-85.0 [46].

Phylogenomic placement of *Colletotrichum* sp. CLE4 was performed using the PHYling_unified (https://github.com/stajichlab/PHYling_unified) pipeline to generate a protein alignment of all species in the final dataset. This pipeline utilizes HMMER v.3 [43] and ClipKIT [100] to search for, build, and trim an alignment based on the BUSCO fungi_odb10 gene set. A maximum likelihood phylogeny was built from this alignment using IQ-TREE2 v.2.2.6 [101], with the -p option to indicate gene partitions [102] and the -m option to run ModelFinder Plus which identifies the optimal evolutionary model for each partition based on BIC [103]. The resulting phylogenetic tree was imported into R and visualized using ggtree v. 3.8.2 [104]. We also used fastANI v. 1.33 to compare the average nucleotide identity of the draft *Colletotrichum* sp. CLE4 genome to the genome of its closest sister taxa based on the whole genome phylogeny.

Phylogenetic hierarchical orthogroups (HOGs) were identified using OrthoFinder v. 2.5.4 [105]. We focused analyses on the phylogenetic node containing all *Colletotrichum* spp. and then compared orthogroup detection and frequency between *Colletotrichum* sp. CLE4 and other members of the *C. actuatum* clade. HOGs were visualized in R using the UpSetR v. 1.4.0 [106] and pheatmap v. 1.0.12 packages [107].

## Results

### Genome structure, repeat landscape, and functional potential of Colletotrichum sp. CLE4

The genome of *Colletotrichum* sp. CLE4 was 48.03 Mbp in total length with 408x coverage, distributed across 168 contigs with an N50 of 506,655 bp and an L50 of 32, indicating a relatively contiguous assembly (Table 1). The genome is haploid, with a predicted genome size based on k-mer frequency profiles of 48.3 Mbp (Figure 1B). BUSCO estimates for the genome using the fungi_odb10 dataset reveal that it is highly complete with 98.8% of the expected single-copy orthologs present and complete. Only 0.3% of the BUSCO genes were fragmented, and 0.9% were missing. Based on these results, we believe that the genome of *Colletotrichum* sp. CLE4 represents a high-quality resource for understanding the ecology and evolution of this isolate.

**Table 1.** Genome and annotation statistics for *Colletotrichum* sp. CLE4. Here we report various AAFTF assembly and Funannotate annotation statistics including the number of contigs in the assembly, the number of contigs of various lengths, the total assembly length, percent GC, N50, L50, number of gene models, percent of genes annotated with different databases, and the number of secreted genes with EffectorP predictions. We also report here the results of the BUSCO assessment using the fungi_odb10 gene set.

| Software | Statistic | *Colletotrichum* sp. CLE4 |
| --- | --- | --- |
| AAFTF | # contigs ($\geq 0$ bp) | 168 |
| | # contigs ($\geq 50,000$ bp) | 127 |
| | Total length ($\geq 0$ bp) | 48,026,924 |
| | Total length ($\geq 50,000$ bp) | 47,411,477 |
|  | Largest contig (bp) | 1,779,047 |
|  | Total length (bp) | 48,026,924 |
|  | Repetitive regions (%) | 2.87 |
|  | GC (%) | 52.29 |
|  | N50 | 506,655 |
|  | L50 | 32 |
|  | # N's per 100 kbp | 5.01 |
| BUSCO (fungi_odb10) | Complete BUSCOs | 98.8 |
|  | Complete and single-copy BUSCOs | 98.4 |
|  | Complete and duplicated BUSCOs | 0.4 |
|  | Fragmented BUSCOs | 0.3 |
|  | Missing BUSCOs | 0.9 |
|  | Total BUSCO groups searched | 758 |
| Funannotate | Total number of gene models | 12,015 |
|  | Number of mRNA genes | 11,678 |
|  | Number of tRNA genes | 337 |
|  | Genes with GO Term (%) | 57.42 |
|  | Genes with InterProScan hit (%) | 78 |
|  | Genes with EggNog hit (%) | 95.53 |
|  | Genes with PFAM hit (%) | 69.13 |
|  | Genes with CAZyme hits (%) | 5.68 |
|  | Genes with MEROPS hits (%) | 3.67 |
|  | Genes with secretion prediction (%) | 11.97 |
| EffectorP | Number of total predicted effectors | 469 |
|  | Number of cytoplasmic effectors | 141 |
|  | Number of apoplastic effectors | 328 |

Repeat content in the genome was relatively low, representing only 2.87% of the total genome, with LTR and unknown elements being most prevalent (Figure 1C). The genome repeat landscape indicates that elements have accumulated gradually through time in this species and also exposes a possible historical expansion of repeat content, corresponding to ∼11-12% divergence (Figure 1D). While similar patterns of LTR and unknown element expansion have been observed in *Colletotrichum* spp., the overall percent repeat content here is less than what has been reported in other species (e.g., 6.08% in *C. truncatum* [108], 5.86% in *C. incanum* [78]).

Annotation of the *Colletotrichum* sp. CLE4 genome using Funannotate identified a total of 12,015 gene models, including 11,678 mRNA genes and 337 tRNA genes, with 95.53% of gene models having EggNog database annotation hits (Table 1). Additionally, 683 (5.68%) genes were identified as having CAZyme domains (Figure S1). Approximately 11.97% of the genes were predicted to encode secreted proteins, including 469 predicted effectors, of which 141 were predicted to be cytoplasmic effectors and 328 were predicted to be apoplastic effectors.

Based on CAZyme content, CATAStrophy predicted that *Colletotrichum* sp. CLE4 was most likely a hemibiotroph, specifically an extracellular (non-appressorial) mesotroph. This classification is described as representing facultative biotrophic species that have longer latent periods than necrotrophs and that invade extracellular host tissues [56]. This classification group includes members that grow biotrophically under optimal environmental conditions, but under variable conditions can cause disease [109,110]. Thus, this assignment is consistent with potential for an opportunistic pathogenic lifestyle.

### Whole-genome phylogenetic placement of Colletotrichum sp. CLE4 and genomic similarity to close relatives

Whole genome phylogenetic approaches place *Colletotrichum* sp. CLE4 in the *C. acutatum* complex (Figure 2), whose common ancestor was dated at 14.5 mya [6]. Within this complex, *Colletotrichum* sp. CLE4 is placed sister to *C. godetiae,* which is best known for causing disease in terrestrial eudicot plants [5,7]. Average nucleotide identity (ANI) between *Colletotrichum* sp. CLE4 and its sister *C. godetiae* was relatively high at 98.98%. While ANI to other members of its immediate clade were lower at 93.50% for *C. salcis* and 93.77% for *C. phormii*. Although ANI species boundaries in fungi have yet to be used extensively to delineate species boundaries, with such high similarity it’s possible that *Colletotrichum* sp. CLE4 may represent a new marine strain of *C. godetiae* that infects monocot plants.

**Figure 2.**
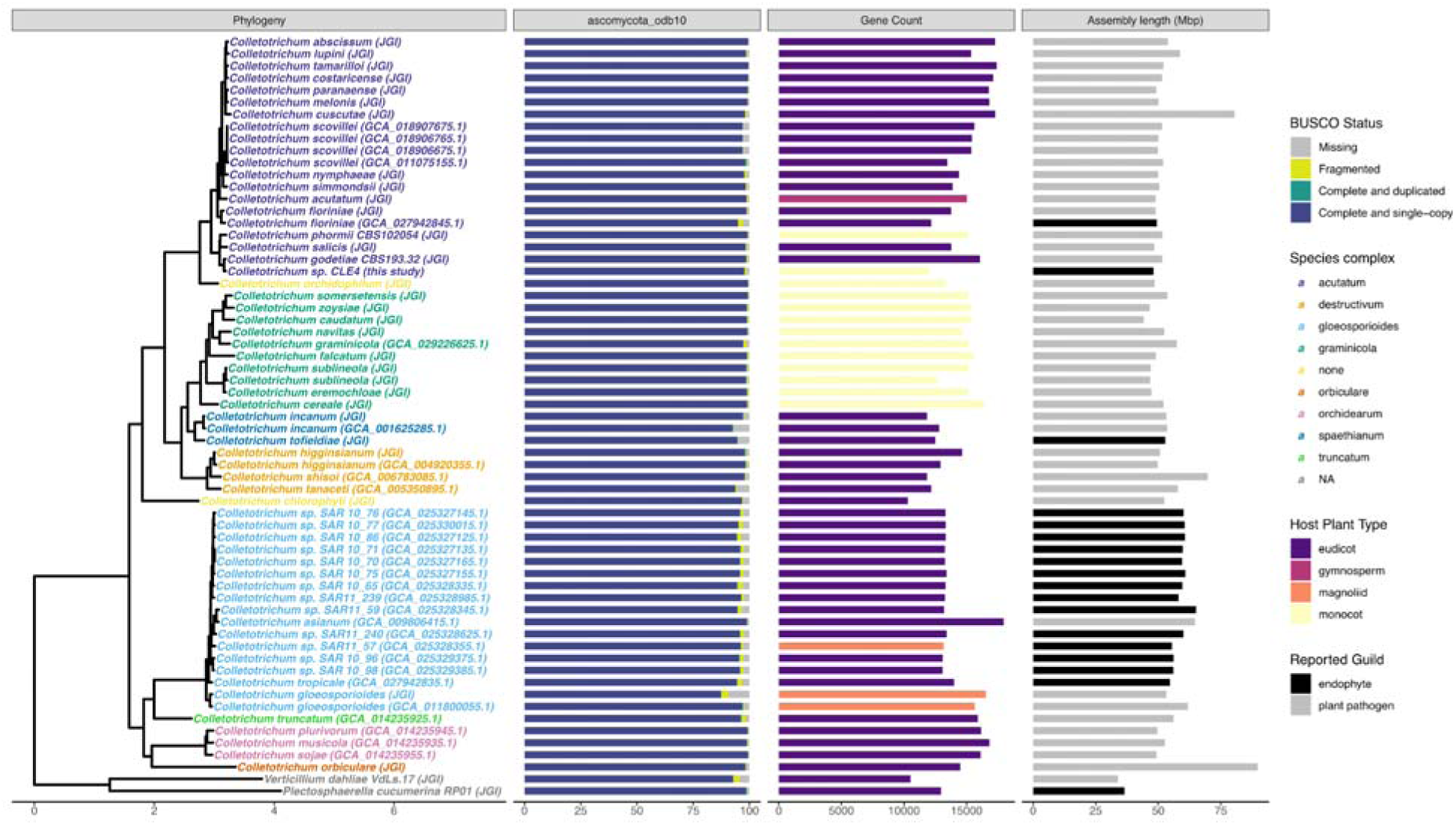
Phylogenetic placement and genomic content comparison to other *Colletotrichum* spp. From left to right, first, a maximum likelihood phylogeny that shows the relationship of *Colletotrichum* sp. CLE4 to other *Colletotrichum* spp. lineages. This tree was generated using IQ-TREE2 on an alignment of BUSCO fungi_odb10 HMMs constructed using the PHYling_unified pipeline. Taxon labels in the phylogeny are shown colored by their assigned *Colletotrichum* species complex. Next (from left to right), in association with this phylogeny, a bar chart of BUSCO “protein” completion status for the ascomycota_odb10 set is shown. Bars show the percentage of genes found in each genome annotation as a percentage of the total gene set and are colored by BUSCO status (missing = gray, fragmented = yellow, complete and duplicated = green, complete and single copy = blue). Next is a bar chart of predicted gene counts for each taxon with counts colored by fungal host plant ecotype reported during isolate deposition (eudicot = purple, gymnosperm = pink, magnoliid = orange, monocot = yellow). Finally, there is a bar chart of the draft genome size (Mbp) for each taxon with genome size colored by fungal guild (endophyte = black, plant pathogen = grey).

*Colletotrichum* sp. CLE4 has the smallest reported genome size and fewest number of gene models of any of the members in the *C. acutatum* complex looked at in this study. The genome size of *Colletotrichum* sp. CLE4 is 8.69% smaller than the average genome size in the *C. acutatum* complex (52.6 Mbp). While the gene content is 21.90% less in comparison to the average number of genes in the complex (15,383). Further, despite high ANI similarity*, Colletotrichum* sp. CLE4 has a genome size that is 7.02% smaller compared to the genome of its sister taxa *C. godetiae* (51.7 Mbp) and has 25.24% less gene content (16,071). Interestingly, the *Colletotrichum* sp. CLE4 genome is only slightly smaller than the only reported endophyte isolate in this complex, *C. fioriniae* [83], with only a 2.86% smaller genome size (49.4 Mbp in *C. fioriniae*) and a 1.34% smaller gene content (12,178 in *C. fioriniae*). Zooming out to the genus overall, while *Colletotrichum* sp. CLE4 is still among the smallest assemblies for genome size and number of gene models, it is not the smallest for either metric. Additionally, the average BUSCO completeness for the *C. acutatum* complex was 98.38% and for the *Colletotrichum* genus was 97.39%. Thus, while smaller in genome size and gene content, the draft genome assembly for *Colletotrichum* sp. CLE4 has a similarly high completion rate (98.8%).

### Gene family reductions in Colletotrichum sp. CLE4

To further explore gene family gain or loss in *Colletotrichum* sp. CLE4, we performed OrthoFinder analysis, comparing *Colletotrichum* sp. CLE4 with other members of the genus *Colletotrichum* with a focus on comparisons to members of the *C. acutatum* complex. In total, we identified 38,833 phylogenetically hierarchical orthogroups (HOGs) among all *Colletotrichum* spp. and 23,298 HOGs among members of the *C. acutatum* complex.

Of these, 9197 HOGs were conserved across all members of the *C. acutatum* complex and *Colletotrichum* sp. CLE4 (Figure 3A). Interestingly, *Colletotrichum* sp. CLE4 appears to be missing 591 HOGs that are shared between all other members of the *C. acutatum* complex, which we infer to be most likely due to gene loss in CLE4. The main functional domains of the missing HOGs include hypothetical domains, transcription factors, transporters, cytochrome p450s, FAD-binding domains, and heterokaryon incompatibility protein domains (Figure 3B). This set also included a handful of HOGs with CAZyme domains including GH3, GH43, GH76, GT25, GT15, and GT90.

**Figure 3.**
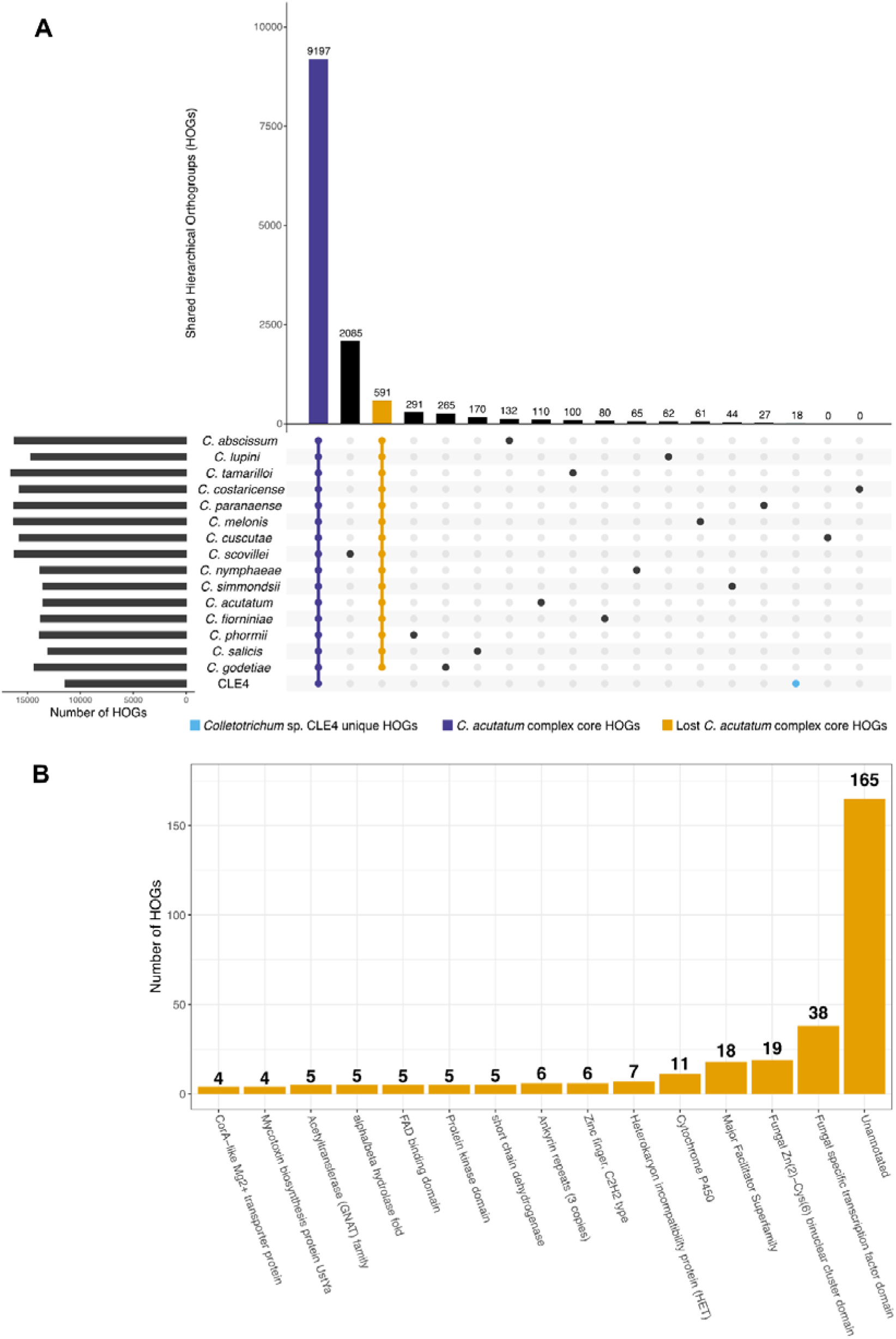
*Colletotrichum* sp. CLE4 is missing many *C. acutatum* complex conserved phylogenetically hierarchical orthogroups (HOGs). (A) UpSet plot depicting shared HOGs within the *C. acutatum* complex with *Colletotrichum* spp. organized by their phylogenetic relationship from Figure 2. Vertical bars display the counts of HOGs in each group while the dots indicate the organism present in each group. Horizontal bars display the number of HOGs in the genome of each organism. We have highlighted three groups of HOGs: unique HOGs to *Colletotrichum* sp. CLE4 (blue), conserved HOGs shared by the *C. acutatum* complex (purple), and conserved HOGs missing in *Colletotrichum* sp. CLE4, but shared by the rest of the *C. acutatum* complex (yellow). (B) Bar plot depicting the distribution of PFAM annotations across the conserved HOGs missing in *Colletotrichum* sp. CLE4 but shared by the rest of the *C. acutatum* complex. To simplify visualization, only annotations that were observed across ≥ 4 HOGs are depicted.

In total, only 18 HOGs were exclusively present in *Colletotrichum* sp. CLE4 and absent in other members of the *C. acutatum* complex, many of which represented uncharacterized proteins (Figure 4). However, only two of these HOGs were exclusively present in *Colletotrichum* sp. CLE4 relative to all other *Colletotrichum* spp, an uncharacterized protein and a short-chain dehydrogenase (Figure S2). Seven HOGs were detected as shared with multiple members of the *C*. *gloeosporioides* complex, which includes many reported endophytic species. Two HOGs were predicted to be apoplastic plant effectors, representing a multicopper oxidase and a FAD-binding domain protein. Further and of particular note, one HOG was predicted to be a NodB homology domain-containing protein.

**Figure 4.**
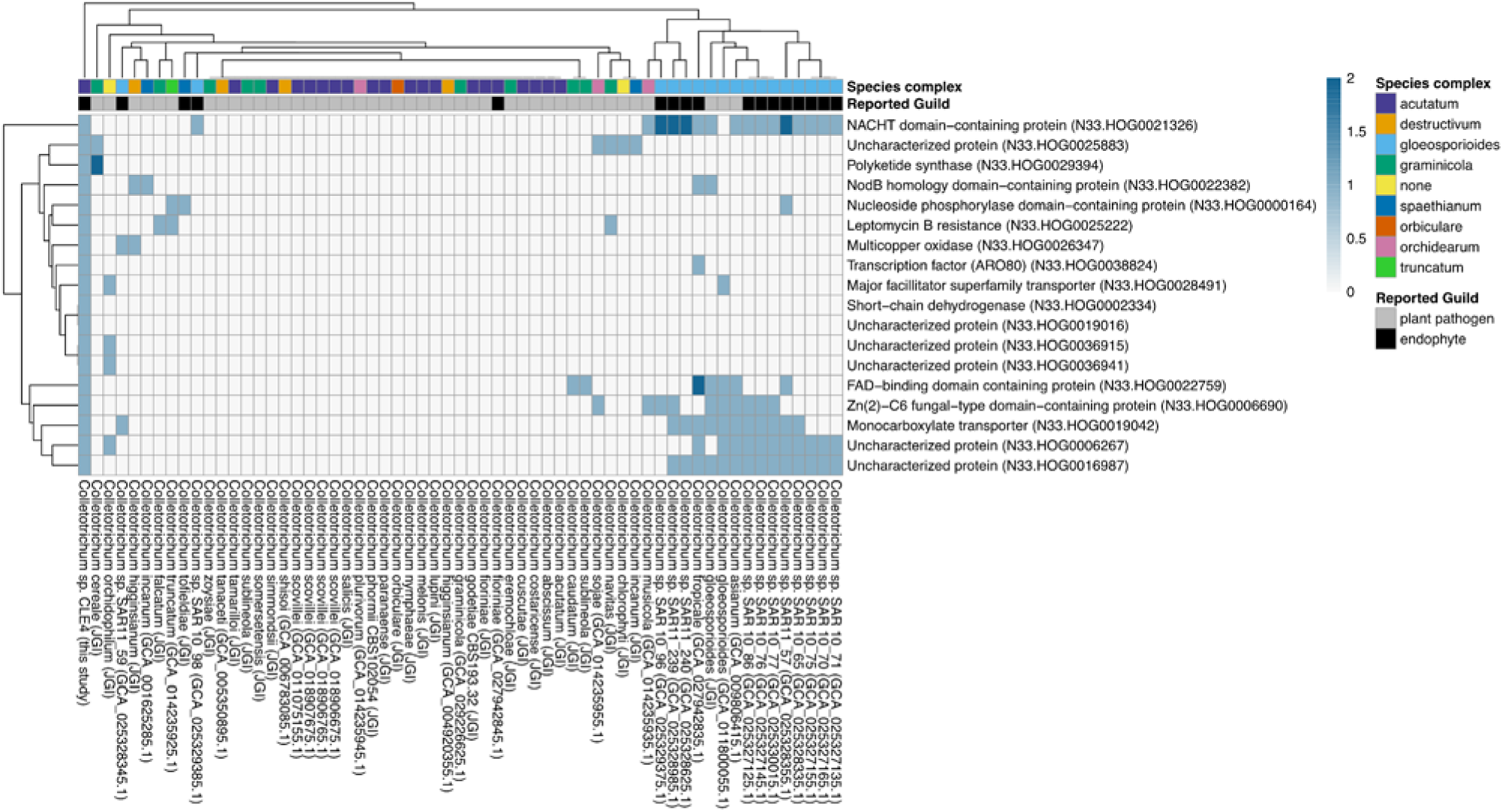
Majority of unique HOGs shared with endophytic *C. gloeosporiodes* complex. A heatmap is depicted here visualizing the copy number of phylogenetically hierarchical orthogroups (HOGs) that were unique to *Colletotrichum* sp. CLE4 relative to other members of the *C. actutatum* complex. Further, the dendrogram to the left of the heatmap clusters the HOGs by similarity in counts between the different HOGs, while the dendrogram above clusters genomes based on similarity in counts between the different genomes.

## Discussion

### Genome and annotation provide critical resource for marine-plant fungi work

This study provides the first draft genome and annotation of a marine *Colletotrichum* sp. and is a valuable resource for future investigations into the evolution and ecology of this highly adaptable fungal genus. At 98.8% BUSCO completion, this genome is comparable to other members of the *C. acutatum* complex and the broader *Colletotrichum* genus. This high level of completeness indicates that the assembly accurately reflects the genetic content of *Colletotrichum* sp. CLE4 and can serve as a foundation for understanding the genomic basis of its adaptation to a marine environment.

Interestingly, the repeat content of the *Colletotrichum* sp. CLE4 genome (2.87%) is lower than that reported for other *Colletotrichum* species [78,108,111]. This could be due to the use of short-read sequencing, which may collapse repetitive regions, thereby underestimating repeat content. However, another study has reported a significant positive correlation between genome size and repeat content in *Colletotrichum* spp. [111]. Further, in *C. tanaceti*, higher repeat content was suggested as a possible mechanism for expansion of pathogenicity genes [89]. Further studies using long-read sequencing could help clarify the repeat landscape of *Colletotrichum* sp. CLE4 and explore its potential role in genomic streamlining and adaptation.

### Phylogenetic placement indicates likely recent evolutionary association

Whole genome phylogenetic approaches place this isolate it in *C. acutatum* complex, which evolved 14.5 mya [6], and place it sister to *C. godetiae,* which is best known for causing disease in a broad range of terrestrial eudicot plants and having a global distribution [5,7]. Interestingly in multi-locus phylogenies, *C. godetiae* places sister to *C. lauri* (no publicly available genome) which has been reported once in association with neither a monocot or eudicot, but instead a magnoliid plant [7,112]. The evolutionary history of the *Colletotrichum* genus suggests an ancestral association with eudicot hosts, with subsequent diversification and independent adaptation to monocot hosts as flowering plants diversified [6]. Seagrasses, such as *Z. marina*, are early branching monocots whose ancestors recolonized the marine ecosystem 70 - 100 mya [113]. The more recent evolution of the *C. acutatum* complex suggests that *Colletotrichum* sp. CLE4’s relationship with seagrass likely occurred after the return of the ancestor of *Z. marina* to the ocean, as opposed to co-evolving with *Z. marina*. This is similar to the timing of other monocot-host jumps in *C. orchidophilum* and *C. phormii*, which both are predicted to have transitioned at a date after the speciation of their host [6]. Ultimately, this means that the ancestor of *Colletotrichum* sp. CLE4 needed to adapt to both a monocot host and the marine ecosystem simultaneously.

Adaptations by seagrasses to the marine ecosystem may pose additional challenges as well. For example, seagrasses have lost all the required genes to form stomata [114], and their cell wall contains polyanionic, low-methylated pectins and sulfated galactans, in addition the polysaccharides typical of land plants [115]. The plant cell wall is considered the first line of plant defense against microbial invasion and the stomata are often how fungi initially invade plants [116,117]. These physiological modifications, as a result of adaptation to the marine ecosystem, would likely make it harder for *Colletotrichum* sp. CLE4 to invade and proliferate in *Z. marina*, possibly requiring a new or divergent invasion strategy relative to the strategies and genes utilized by close relatives for land plant colonization.

### Genomic streamlining during adaptation to a marine monocot host

Comparative genomic analyses revealed smaller genome size, gene count, and absence of conserved gene families (i.e., HOGs) in *Colletotrichum* sp. CLE4 compared to other members of the *C. acutatum* complex. Specifically, we found that the genome size of *Colletotrichum* sp. CLE4 was 8.69% smaller, and its gene content was 21.90% lower than the average for the *C. acutatum* complex. This combined with the absence of 591 gene families that are conserved among all other *C acutatum* complex members leads us to conclude that the smaller genome, gene count and reduced orthologous groups may be the result of genome reduction and streamlining. Genome streamlining is a well-documented phenomenon in microbial adaptation to marine environments, as seen in bacteria [118], and has been recently described in fungi [119]. Streamlining removes non-essential genes and non-coding DNA to improve efficiency, often at the cost of metabolic versatility. For example, the marine fungus *Rhodotorula sphaerocarpa* exhibits a 10% smaller genome size compared to its terrestrial relatives, largely due to a decrease in transporter genes, particularly Major Facilitator Superfamily transporters, which are key for cross-membrane transport of organic solutes [119].

Genomic streamlining in *Colletotrichum* sp. CLE4 could also be linked to adaptation to a monocot host or an endophytic lifestyle. Pathogens often have expanded or unique secreted enzymes related to host-specialization and virulence in comparison to non-pathogens, and similar patterns have been reported for hemibiotrophic fungi compared to biotrophs [2,92,120–122]. However, gene family differences in some studies have been reported to be more about relatedness than trophic lifestyle [123]. Previous work in *Colletotrichum* species have observed gains and losses of CAZyme and protease encoding genes in species that have a more narrow host range and have suggested that switching to a new host involves gene losses coupled with expansions in lineage-specific genes [6,61,78,124].

*Colletotrichum* sp. CLE4 had 683 CAZyme domain predictions, which is lower than most other *Colletotrichum* species. Monocot infecting *Colletotrichum* spp. are generally reported to have a smaller number of CAZymes compared to eudicot infecting species (741 vs. 867 on average) [6]. This supports that some gene content reduction may be due to specialization to a monocot host. However, *Colletotrichum* sp. CLE4 has a 7.82% smaller CAZyme content when compared to the average for other monocot infecting species, indicating that host specialization alone may not fully explain the extent of its smaller gene content.

### Retention and loss of gene families provides functional insight into adaptation

In comparison to other *C. acutatum* complex members, *Colletotrichum* sp. CLE4 has lost 591 conserved gene families. While the majority of these had no predicted function, there was an enrichment in the loss of transcription factors, transporters, cytochrome p450s, and FAD-binding domains, as well as some specific CAZymes. The loss of gene families with transporter domain annotations, particularly Major Facilitator Superfamily transporters, is similar to the reports from genomic streamlining in marine *Rhodotorula* in response to marine adaptation [119].

While the reduction in transporters may relate to adaptation to the marine realm, the loss of other gene families, such as cytochrome P450s and CAZymes, could relate to specialization to a monocot host or endophytic lifestyle. It’s been suggested that having a diverse set of cytochrome p450s play a role in the colonization success or virulence of plant pathogenic fungi [122,125]. In *Colletotrichum* spp., contractions in cytochrome p450 diversity have been suggested to relate to host range and specificity [6,61,126,127].

Similarly, the loss of certain CAZyme gene families further highlights the specialization of *Colletotrichum* sp. CLE4, potentially reflecting adaptations not just to marine life but also to the unique defenses of its monocot host and lifestyle strategy. For example, GH3, which can help detoxify plant antifungal commands, may be unnecessary depending on the specific defenses of *Z. marina* [128]. GH43 has been suggested to be important for plant-host interaction or plant tissue degradation in other *Colletotrichum* spp. [6] and expanded across distantly related pathogenic lineages [78]. While GH76 expansions in *Colletotrichum* have been associated with host-specificity towards woody plants [124].

A study comparing gene family differences between monocot and eudicot-infecting *Colletotrichum* species found that pathogenic eudicot species retained three unique gene families that were lost in monocot-infecting species, including a secreted β-glucosidase (GH3); a secreted protein with a FAD-binding domain, and an α-1,2-mannosidase (GH92) [6]. The loss of GH3 and FAD-binding domains in *Colletotrichum* sp. CLE4 aligns with this pattern, suggesting that these losses may relate to specialization to a monocot host.

In comparison to other *C. acutatum* complex members, *Colletotrichum* sp. CLE4 had only 18 unique gene families, of which only two were truly unique, while several others were shared with endophytic members of *C. gloeosporioides* species complex. Two gene families were predicted to be apoplastic plant effectors, representing a multicopper oxidase and a FAD-binding domain protein. While often associated with pathogenicity, effectors are also important for beneficial plant-fungal interactions [129–132] and for biotrophic lifestyles, where fungi may still need to suppress host defenses and evade recognition [133,134]. Further and of particular note, one gene family was predicted to be a NodB homology domain-containing protein. NodB genes are chitin deacetylases involved in the production of signaling molecules, most famously in legume-rhizobia symbiosis [135,136], and homologous signaling pathways have been used by symbiotic fungi [137]. The retention of gene families shared with *C. gloeosporioides* endophytes aligns with previous studies suggesting that such shared gene families across distant *Colletotrichum* species result from recent independent acquisitions or rapid losses during host specialization [2,6,61,78].

### Pathogen or endophyte - deconvoluting a complex hemibiotrophic lifestyle

*Colletotrichum* spp. isolated from healthy, undamaged seagrass tissues, such as *Zostera marina*, have not been associated with anthracnose or other known pathogenic symptoms in marine plants, suggesting they may act primarily as endophytes in these environments. However, given the well-documented pathogenic potential of *Colletotrichum* spp. in terrestrial plants and their hemibiotrophic lifestyle, we used machine learning with CATAStrophy to predict the lifestyle of *Colletotrichum* sp. CLE4.

Perhaps unsurprisingly, CATAStrophy predicted that *Colletotrichum* sp. CLE4 is a hemibiotroph, specifically an extracellular mesotroph. This lifestyle involves a biotrophic phase, where the fungus maintains the host plant’s viability under favorable conditions, followed by a necrotrophic phase under stress, where it acts as a pathogen. This prediction aligns with CATAStrophy’s classification of most other *Colletotrichum* spp. as mesotrophs. This dual capacity for both benign and pathogenic behavior is similar to fungi like *Cladosporium fulvum*, which exhibits a biotrophic lifestyle under controlled, optimal conditions but can turn pathogenic in response to environmental stress [109,110].

The predicted potential of *Colletotrichum* sp. CLE4 to switch between endophytic and pathogenic roles suggests that it could remain a benign endophyte under stable conditions but become harmful under environmental stressors or when *Z. marina* is compromised. This flexibility supports a growing view of many fungi as multi-niche organisms that can form either beneficial or pathogenic associations depending on context [138]. Thus, while *Colletotrichum* sp. CLE4 likely functions as an endophyte in *Z. marina*, it may retain the genomic potential to become pathogenic under adverse conditions, highlighting the importance of environmental factors in shaping fungal-host dynamics, and a need for further work in understanding the exact nature of its ecological role when associated with *Z. marina*.

## Conclusion

We report the first high quality draft genome and annotation of a marine monocot infecting *Colletotrichum* sp. The genome is near-complete (98.8%) and encodes 12,015 genes, of which 5.7% are CAZymes and 12.6% are predicted secreted proteins. Whole-genome phylogenetic analyses place *Colletotrichum* sp. CLE4 within the *C. acutatum* complex, most closely related to *C. godetiae*, which infects terrestrial eudicot plants. Overall, we found evidence of a streamlined genome, with an 8.69% reduction in genome size, 21.90% reduction in gene content, and a loss of 591 conserved gene families compared to other members of the *C. acutatum* complex. This streamlining is likely due to adaptation to both the marine ecosystem and a monocot host. We also identified unique gene families some of which were shared with members of the *C*. *gloeosporioides* complex which includes several endophytes, as well as NodB homology containing domain protein. Machine learning analyses predicted that *Colletotrichum* sp. CLE4 has an extracellular mesotroph lifestyle, which may indicate it still has capacity to serve as an opportunistic pathogen of *Z. marina*. This study provides a foundational insight into understanding the evolutionary trajectory and ecological adaptations of marine-plant associated *Colletotrichum* spp. Further work is needed, including challenge experiments and transcriptomics, to assess whether CLE4 is endophyte of ZM only under optimal conditions and whether new environmental stressors such as a changing climate might trigger opportunistic pathogenicity.

## DNA Deposition

The raw sequences were deposited at GenBank under accession no. <u>PRJNA1140278</u>. This Whole Genome Shotgun project has been deposited at DDBJ/ENA/GenBank under the accession JBFUUI000000000. The version described in this paper is version JBFUUI010000000. All code used in this work has been deposited on Github (<u>casett/ZM_Colletotrichum_sp_Genome</u>) and archived in Zenodo (DOI: <u>10.5281/zenodo.14207532</u>).

## Author Contributions

CLE conceived and designed the experiments, performed sampling, analyzed the data, prepared figures and/or tables, wrote and reviewed drafts of the paper. JAE and JES reviewed drafts of the paper.

## Acknowledgements

We would like to thank the following undergraduate researchers for their contributions to this work including help with sample collection, fungal isolation and DNA extraction: Tess McDaniel, Katelin Jones, Katie Somers and Neil Brahmbhatt. We would also like to thank John J. Stachowicz for use of his scientific sampling permit, California Department of Fish and Wildlife Scientific Collecting Permit # SC 4874.

