## Supplemental Figures and Table Legends for "Genomic streamlining of seagrass-associated *Colletotrichum* sp. may be related to its adaptation to a marine monocot host"

**Supplemental Table Legends and Figures:**

**Table S1.** Genomes used in phylogenetic and orthogroup analyses of *Colletotrichum* sp. CLE4. Here we provide details on each genome obtained from NCBI or JGI including taxonomic information, host type and ecosystem reported upon collection, predicted ecological guild, assembly statistics, BioProject number, assembly accession number, and appropriate data citation.

**Figure S1.** Effector and CAZyme profiles for *Colletotrichum* sp. CLE4. (A) Bar plot depicting the number of secreted genes in the genome that are predicted to be plant apoplastic and cytoplasmic effectors. (B) Bar plot depicting the frequency distribution of predicted CAZymes in the genome across CAZyme families.

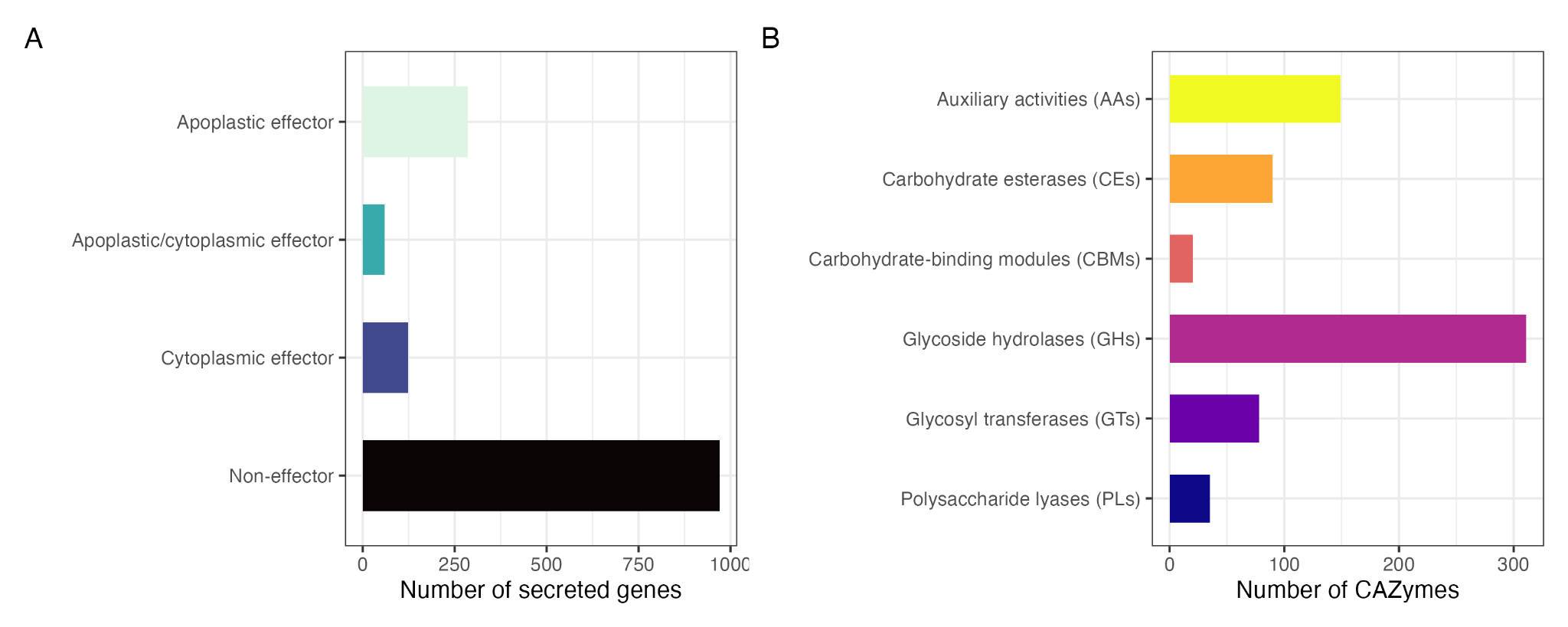

**Figure S2.** *Colletotrichum* sp. CLE4 is missing many *Colletotrichum* species conserved HOGs. (A) UpSet plot depicting shared HOGs across *Colletotrichum* species complexes. Vertical bars display the detection of HOGs in each group while the dots indicate the complex or organism present in each group. Horizontal bars display the number of HOGs detected in the complex or organism. We have highlighted three groups of HOGs: unique HOGs to *Colletotrichum* sp. CLE4 (blue), conserved HOGs shared by all *Colletotrichum* species (purple), conserved HOGs missing in *Colletotrichum* sp. CLE4, but shared by the rest of *Colletotrichum* species (yellow), and conserved HOGs between *Colletotrichum* sp. CLE4 and the *C. acutatum* complex (orange).

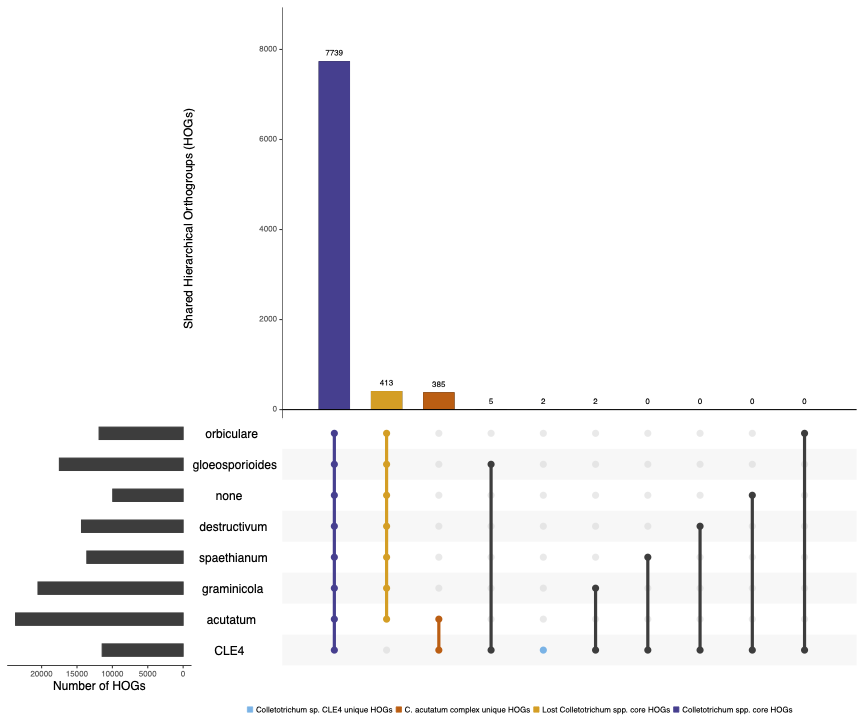
